# Characterization of PROTAC specificity and endogenous protein interactomes using ProtacID

**DOI:** 10.1101/2025.04.14.648778

**Authors:** Suman Shrestha, Matthew ER Maitland, Laili Jing, Shili Duan, David Y. Nie, Jonathan St-Germain, Dalia Barsyte-Lovejoy, Cheryl H Arrowsmith, Brian Raught

## Abstract

Here we describe ProtacID, a flexible BioID (proximity-dependent biotinylation)-based approach to identify PROTAC-proximal proteins in living cells. ProtacID analysis of VHL- and CRBN-recruiting PROTACs targeting a number of different proteins (localized to chromatin or cellular membranes, and tested across six different human cell lines) demonstrates how this technique can be used to validate PROTAC degradation targets and identify non-productive (*i*.*e*. non-degraded) PROTAC-interacting proteins, addressing a critical need in the field of PROTAC development. We also demonstrate that ProtacID can be used to characterize native, endogenous multiprotein complexes without the use of antibodies, or modification of the protein of interest with epitope tags or biotin ligase tagging.

## Introduction

One of the most exciting and fastest moving areas of modern drug discovery, Targeted Protein Degradation (TPD) harnesses the power of the endogenous ubiquitin proteasome system to specifically degrade proteins linked to disease^1, 2^. PROteolysis TArgeting Chimeras (PROTACs) are bivalent, cell permeable TPD compounds consisting of a “warhead” moiety that interacts with one or more target proteins, and a ubiquitin E3 ligase “recruiter” component, connected via a chemical linker (**Fig. 1a**, inset). PROTACs thus direct an endogenous E3 ligase to “neo-substrates” to effect ubiquitylation, resulting in 26S proteasome-mediated degradation. The majority of the >5000 PROTACs^3^ reported to date utilize just two ubiquitin E3 ligases: (i) VHL (Von Hippel-Lindau), the substrate targeting receptor subunit of a Cullin 2 (CUL2)-RING E3 Ligase complex (CRL2^VHL^) or (ii) CRBN (cereblon), the substrate receptor subunit of a CUL4-based complex (CRL4^CRBN^). PROTACs serve as important research tools, and multiple PROTAC-based drugs have recently entered clinical trials^1^.

**Figure 1.**
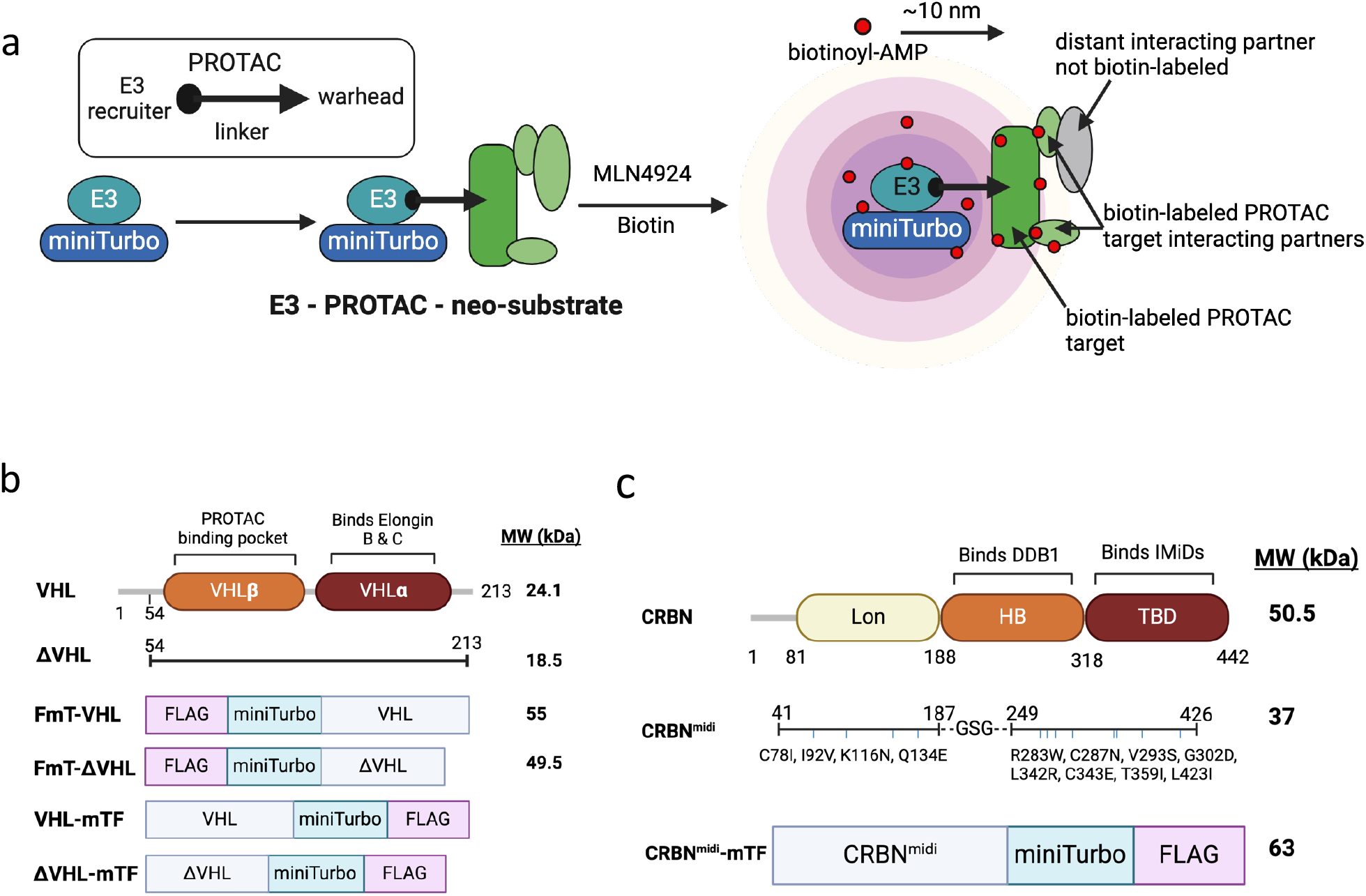
(**a**) A PROTAC molecule consists of “warhead” and “E3 recruiter” moieties connected by a chemical linker (*inset*). ProtacID is conducted by (i) stably expressing a relevant E3 ligase subunit (*e*.*g*. VHL or CRBN) fused with a BioID tool such as miniTurbo^12^. miniTurbo is an abortive biotin ligase that generates chemically reactive biotin (biotinoyl-AMP, *red circles*), which diffuses away from the enzyme and reacts with amine groups on lysine residues in nearby proteins. (ii) In the presence of a PROTAC, neo-substrates are recruited to the E3-miniTurbo protein - but are not ubiquitylated and degraded in the presence of the neddylation inhibitor MLN4924. Lysine residues in proteins within ∼10 nm (an average protein has a Stokes radius of 4-5 nm) are susceptible to biotinylation. (iii) Following cell lysis, biotinylated proteins are isolated using streptavidin-sepharose, and identified using mass spectrometry. (**b**) Schematic diagram of the FLAG-miniTurbo (FmT)-tagged VHL fusion protein variants tested here. (**c**) Schematic of the CRBN^midi^-mTF fusion protein used for ProtacID.

Importantly, however, while PROTAC development has surged in popularity, our ability to characterize PROTAC specificity in living cells has lagged behind. A common method for assessing TPD tool specificity is global proteome analysis, which identifies proteins that decrease in abundance - via so-called “productive interactions” - following PROTAC treatment^4^. While powerful, this approach has some important limitations. The loss of a particular protein in response to PROTAC treatment does not necessarily indicate that it is a direct PROTAC target. For example, if a PROTAC-targeted protein plays an important regulatory role upstream of one or more additional short-lived polypeptide(s), or happens to be a key structural component of a protein complex, its loss can trigger the secondary loss of multiple additional proteins^5-7^. Conversely, PROTAC binding does not always result in the degradation of a protein interactor, yet could still affect its activity, localization or interactions with binding partners. These so-called “non-productive” PROTAC-target interactions are not detected in global proteome analyses.

Proximity-dependent labeling techniques such as BioID are now widely used to identify protein-protein interactions in living cells^8, 9^. As previously demonstrated with CRL4^CRBN^, tagging an E3 ligase with a BioID tool can also be used to effect PROTAC (or molecular glue)-mediated recruitment of a biotin ligase to an endogenous neo-substrate^10^. We reasoned that applying a similar approach to the VHL protein and the new CRBN^midi^ tool (a recently developed stable CRBN protein variant^11^) could be used to: (i) validate PROTAC targets identified in standard global proteome analyses, (ii) identify non-productive PROTAC interactors, and (iii) characterize PROTAC target interacting protein partners (**Fig. 1a**).

## Results

### ProtacID tool development

To conduct “ProtacID”, we fused a FLAG-tagged miniTurbo^12^ (FmT) enzyme to the N-or C-terminus of full length VHL or a VHL variant lacking unstructured residues 1-53 (ΔVHL, as in^13^) (**Fig. 1b**). The resulting fusion proteins were stably expressed (at levels lower than the endogenous VHL protein, **Supplementary Figure 1a**) in human embryonic kidney 293 Flp-In cells. Cells were treated with: (a) the neddylation inhibitor MLN4924 to block cullin activity (and thus rescue CRL2^VHL^ targets), (b) DMSO or the PROTAC ACBI1^14^ and (c) biotin (**Fig. 2a**). Following cell lysis, biotinylated proteins were affinity purified using streptavidin-sepharose beads.

**Figure 2.**
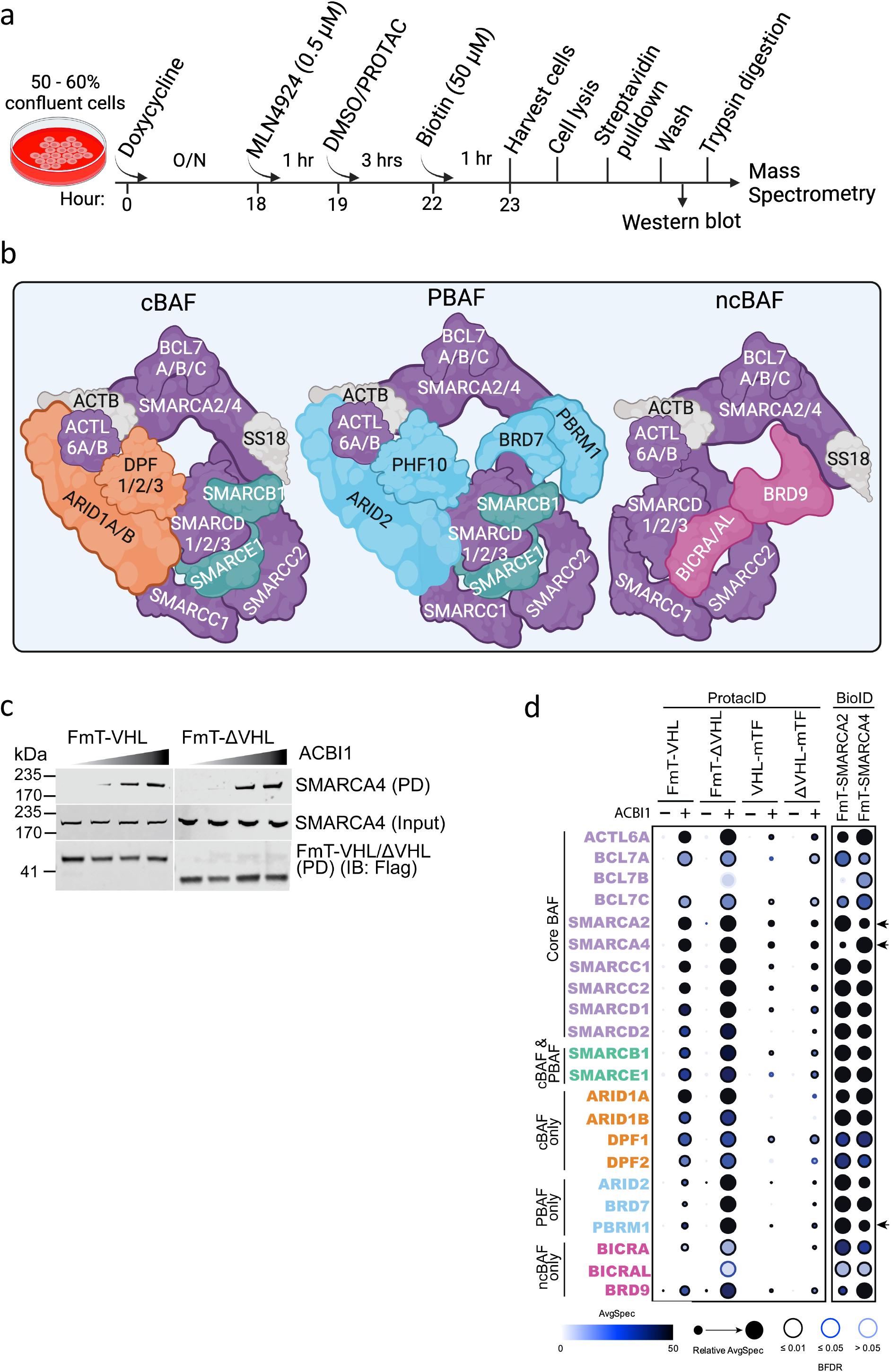
(**a**) ProtacID treatment timeline. See methods for details. (**b**) Schematic diagrams of the canonical (cBAF), non-canonical (ncBAF) and polybromo-associated (PBAF) BAF complex variants. Components specific to each BAF variant are colour-coded as in panel d. (**c**) FmT-VHL proteins form a ternary complex with ACBI1 and SMARCA4. MLN4924 and increasing doses of ACBI1 (0, 10, 100, 1000 nM) were applied to 293 Flp-In cells expressing the indicated FmT-tagged VHL proteins. After biotin treatment, cells were lysed, and biotinylated proteins isolated using streptavidin-sepharose. Isolated proteins (PD, *pulldown*) or whole cell lysates (*input*) were subjected to western blotting with the indicated antibodies. IB, immunoblot. (**d**) FmT-VHL tool testing. ProtacID was conducted on 293 Flp-In cells expressing the indicated FmT-VHL protein variants, and treated with either DMSO (-) or 200 nM ACBI1 (+). The Dot Plot depicts proteins identified by ProtacID, where dot size indicates relative peptide abundance across experiments, dot shade indicates average peptide counts, and dot border indicates protein identification confidence level. Standard BioID (*right*) was also conducted in 293 Flp-In cells expressing FmT-SMARCA2 or FmT-SMARCA4. PROTAC neo-substrates are indicated by arrowheads (*right*). Proteins grouped according to BAF variant.

BAF (BRG1/BRM-associated factor) protein complexes are ∼2MDa multi-subunit SWI/SNF chromatin remodelers^15^. Three distinct BAF complex variants have been described: canonical (cBAF), non-canonical (ncBAF) and polybromo-associated (PBAF)^15^ (**Fig. 2b**). A number of VHL-recruiting PROTACs targeting BAF complex components have been developed, including ACBI1^14^, which targets the bromodomains of the SMARCA2 and SMARCA4 ATPases (interchangeable components of all three BAF complex variants) and PBRM1 (a PBAF-specific protein).

An aliquot (∼10%) of the protein-bound beads was subjected to SDS-PAGE and western blotting analysis. SMARCA4 was isolated (and therefore biotinylated) following ACBI1 treatment (but not in the absence of PROTAC), and SMARCA4 biotinylation increased in a PROTAC dose-dependent manner (**Fig. 2c, Supplementary Fig. 2**). FmT-tagged VHL/ΔVHL can thus form a ternary complex with SMARCA4 and ACBI1 to effect SMARCA4 biotinylation.

The remainder of the beads was treated with trypsin, and the eluted peptides identified using mass spectrometry. To first identify VHL-specific interactors, the DMSO-treated (*i*.*e*. no PROTAC) samples were compared to data from 293 Flp-In cells expressing the FmT protein alone, using the SAINTexpress algorithm^16^. High confidence proximity interactors of FmT-VHL/ΔVHL were defined as those with a Bayesian false discovery rate (BFDR) ≤0.01. As expected, FmT-VHL/ΔVHL interacted specifically with other components of the CRL2^VHL^ complex (CUL2, ELOB/C), the well-characterized VHL substrates HIF1A and ARNT/HIF1B, and other previously reported VHL interactors^17, 18^ (**Supplementary Table 1**).

PROTAC-mediated interactions were defined as proteins detected in PROTAC-treated samples that displayed a ≥2-fold (log_2_) increase in peptide counts over those observed in DMSO-treated samples. As expected, SMARCA2, SMARCA4 and PBRM1 were identified as high confidence interactors following ACBI1 treatment in all four ProtacID experiments (*i*.*e*. full length or ΔVHL, FmT-tagged at the N- or C-terminus; **Fig. 2d, Supplementary Table 1**). Standard BioID^19^ was also conducted using 293 Flp-In cells stably expressing FmT-SMARCA2 or FmT-SMARCA4 (**Fig. 2d, Supplementary Table 2**). ACBI1 ProtacID conducted with the FmT-ΔVHL tool yielded the highest number of high-confidence proximity interactor identifications, comprising 21 of the 22 BAF subunits identified in the standard SMARCA2/4 BioID analyses (**Fig. 2d, Supplementary Table 2**). FmT-ΔVHL was thus used in all subsequent experiments.

### ProtacID can distinguish between BAF complex variants

Like ACBI1, ACBI2^20^ and AU-15330^21^ are VHL-recruiting PROTACs that target the bromodomains of SMARCA2, SMARCA4 and PBRM1. VZ185^22^ is a VHL-based PROTAC directed against the bromodomains of the BRD7 and BRD9 proteins, subunits specific to the PBAF and ncBAF complexes, respectively^15^. ProtacID conducted with ACBI2 and AU-15330 displayed extensive (100% and 85%, respectively) overlap with the ACBI1 dataset (**Fig. 3, Supplementary Table 3**). As expected, ProtacID conducted with VZ185 identified both BRD7 and BRD9, along with additional components of the ncBAF and PBAF complexes (**Fig. 3, Supplementary Table 3**). Importantly, however, cBAF-exclusive subunits (ARID1A/B and DPF1/2) were not identified in the VZ185 ProtacID analysis (**Fig. 3, Supplementary Table 3**). Together, these data demonstrate that ProtacID can be used to validate PROTAC targets and identify proximal interacting protein partners of PROTAC target proteins in living cells, and indicate that ProtacID can distinguish between closely related multiprotein complexes.

**Figure 3.**
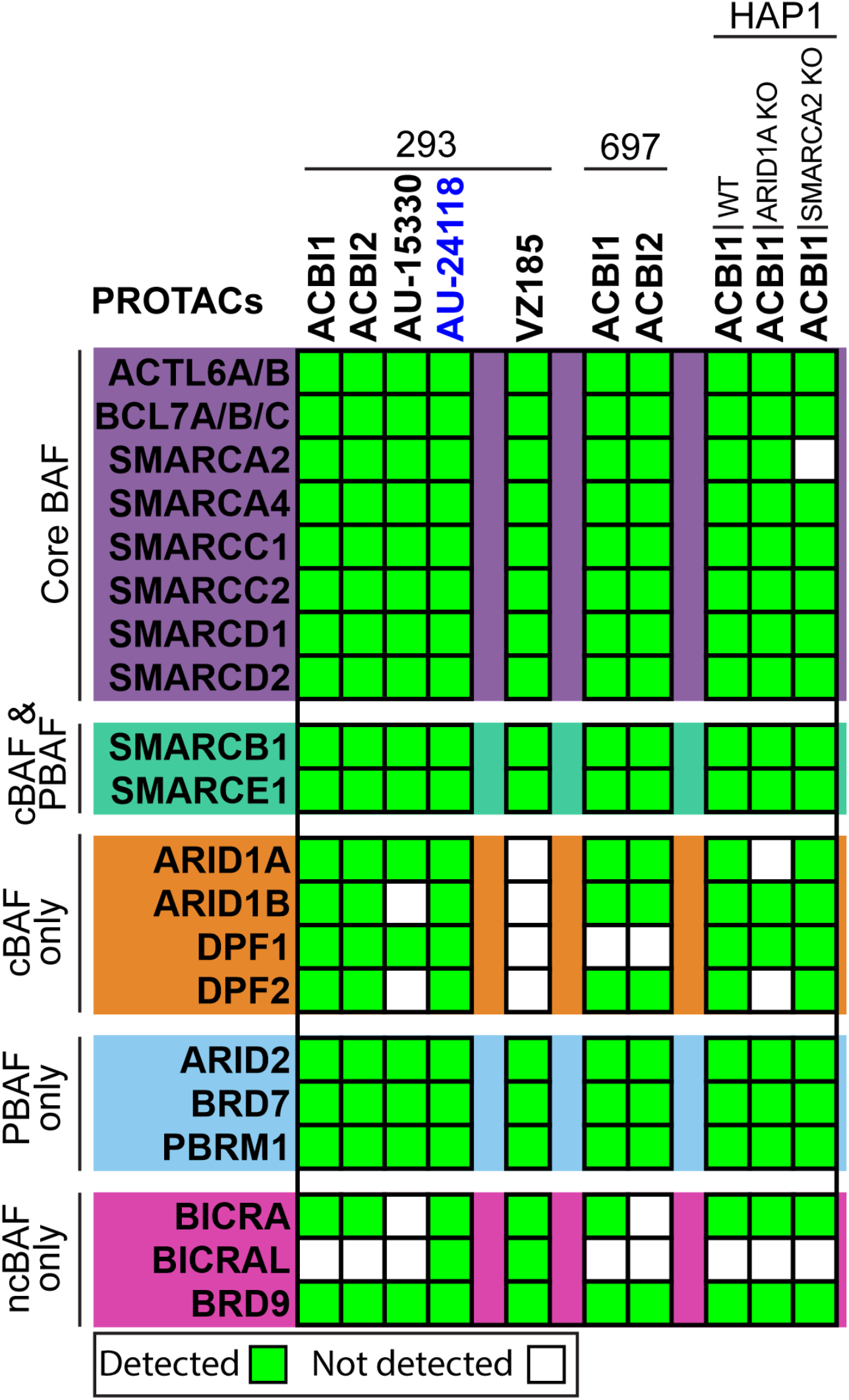
Five different BAF PROTACs were tested across three different human cell lines, as indicated. BAF complex components grouped according to BAF variant. AU-24118 (*blue* font) is a CRBN-recruiting PROTAC, all others are VHL-recruiting PROTACs. *green* - high confidence protein identification. *white* - not identified. Proteins grouped according to BAF variant, as in Fig 2.

### ProtacID can be used across cell lines

ProtacID conducted in FmT-ΔVHL-expressing 697 cells (a human pre-B leukemia line) using ACBI1 and ACBI2 yielded a dataset with >95% overlap with that observed in 293 cells (**Fig. 3, Supplementary Table 4**). ACBI1 ProtacID was also conducted in FmT-ΔVHL-expressing wild type (WT) HAP1 cells (a near-haploid human leukemia cell line), and HAP1 *SMARCA2* and *ARID1A* knockout lines. The WT HAP1 BAF ProtacID dataset was identical to that identified in 293 cells, and HAP1 KO cell datasets specifically lacked the proteins encoded by each *KO* gene (**Fig. 3, Supplementary Table 5**). ProtacID can thus yield reproducible, and highly accurate PROTAC proximity data across cell lines, and in the same cell line with discrete genetic changes.

### CRBN^midi^ ProtacID

The WT CRBN protein is unstable in the absence of its CRL binding partner DDB1^23^, limiting its use as an analytical tool in living cells. CRBN^midi^, a stable CRBN protein variant that displays an extended half-life, was recently developed by the Ciulli lab^11^. N- and C-terminally tagged FmT-CRBN^midi^ proteins were stably expressed in 293 Flp-In cells (**Supplementary Fig 1b**), and tested in ProtacID with the CRBN-recruiting BAF PROTAC AU-24118^24^, which targets SMARCA2/4 and PBRM1 via the same warhead present in ACBI1 and AU-15330. The CRBN^midi^-mTF variant (**Fig. 1c, Supplementary Fig 1b**) yielded the most robust dataset, identifying SMARCA2, SMARCA4, PBRM1 and the same set of BAF complex components identified with VHL-directed BAF PROTACs (**Fig. 3, Supplementary Table 6**). ProtacID can thus be used for the analysis of both VHL- and CRBN-based PROTACs, together comprising the vast majority of current TPD tool compounds.

### ProtacID identifies a non-productive PROTAC-protein interaction

Notably, KIF20B (a kinesin motor protein linked to cell division^25^) was detected as a high confidence interactor in ACBI1 ProtacID conducted with all four FmT-VHL fusion variants in all three cell lines tested here, but was not detected in ACBI2, AU-15330, VZ185 or AU-24118 ProtacIDs. KIF20B was also not detected in standard FmT-SMARCA2 or FmT-SMARCA4 BioID analyses (**Fig. 4a, Supplementary Tables 1-6**). Consistent with this observation, KIF20B was biotinylated in an ACBI1 dose-dependent manner, but did not appear to undergo ternary complex formation in the presence of ACBI2 or AU-15330 (**Fig. 4b** pulldown, PD). Consistent with the original report^14^, our own 293 Flp-In cell global proteome analysis (**Fig. 4c, Supplementary Table 7**) and western blotting (**Fig. 4b** input) indicated that KIF20B is not degraded following ACBI1 treatment. Finally, while SMARCA2, SMARCA4 and PBRM1 were not detected, KIF20B was also detected as an ACBI1 interactor in ProtacID conducted in the absence of MLN4924, via both western blotting and mass spectrometry (**Fig 4d, Supplementary Table 8**), indicating that it is a non-productive ACBI1 interactor (**Fig 4e**). ProtacID can thus also be used to identify non-productive PROTAC-protein interactions.

**Figure 4.**
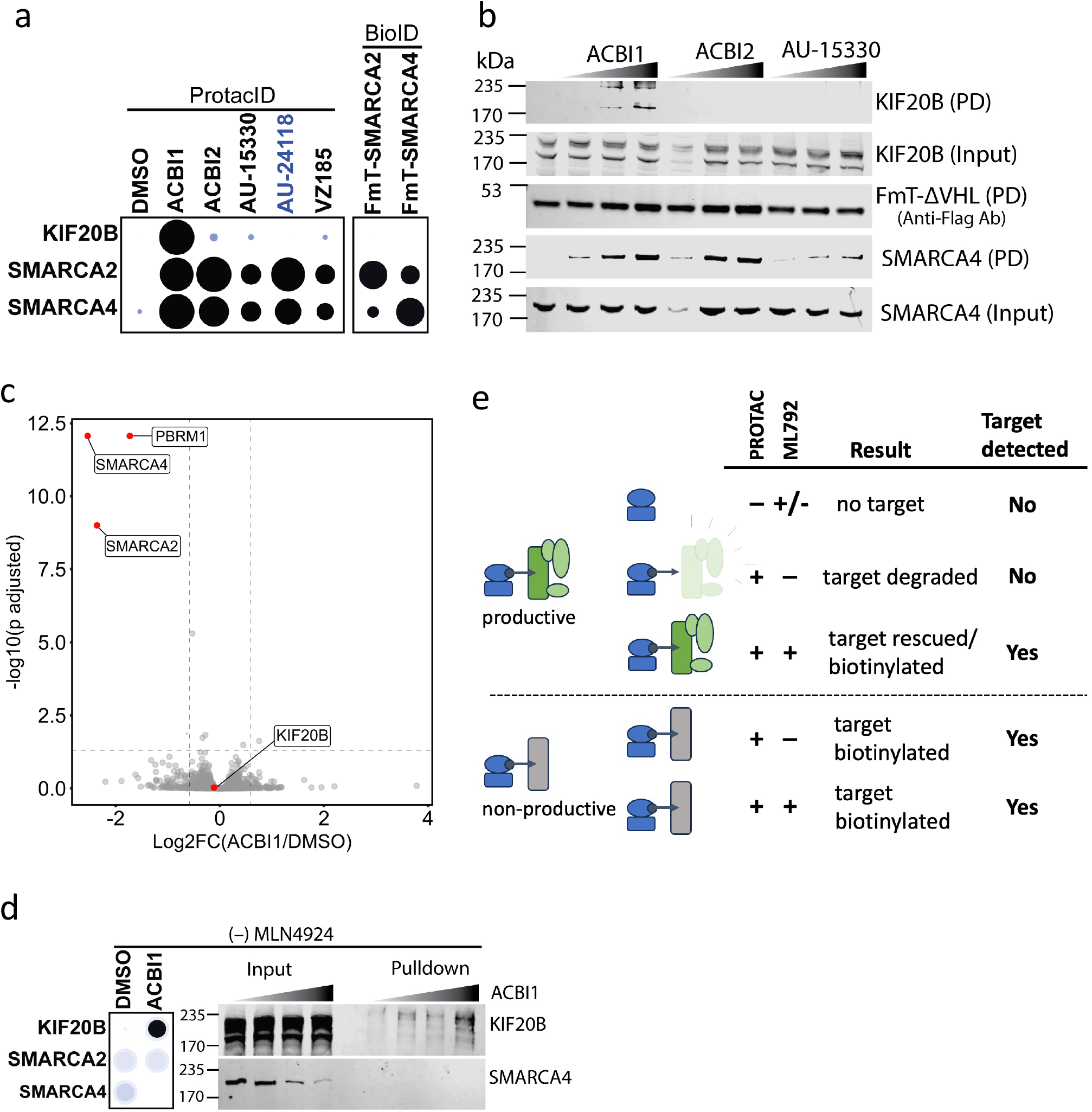
(**a**) Dot Plot highlighting KIF20B, SMARCA2 and SMARCA4 identification in ProtacID analyses (293 cells) with the indicated PROTAC, or standard BioID analysis. (**b**) 293 cells expressing FmT-ΔVHL were treated with increasing concentrations (0, 10, 100, 1000 nM) of the indicated PROTACs, biotin was added, and streptavidin pulldowns conducted as above. Isolated proteins (pullown, PD) or whole cell lysates (*input*) were subjected to western blotting with the indicated antibodies. (**c**) Global proteomics analysis of ACBI1-treated 293 Flp-In cells. Cells were treated for 3 hrs with DMSO or ACBI (1uM) (n = 5 biological replicates). 500ng whole cell lysate was analyzed on a Bruker TIMS-TOF mass spectrometer in DIA mode. See Methods for details. (**d**) (*left*) Dot Plot highlighting KIF20B, SMARCA2 and SMARCA4 identifications in an ACBI1 ProtacID experiment conducted in the absence of the neddylation inhibitor, MLN4924. (*right*) Western blotting results of the same experiment (ACBI1 gradient 0, 10, 100, 1000 nM). (**e**) ProtacID workflow for the identification of productive and non-productive PROTAC-protein interactions. ProtacID conducted in the presence of a PROTAC and MLN4924 can identify both productive and non-productive PROTAC interactors, and their interacting protein partners. ProtacID conducted in the presence of a PROTAC, but in the absence of MLN4924, will identify only non-productive PROTAC interactors.

### ProtacID characterization of BET-targeting PROTACs

Bromodomain and extraterminal domain (BET) family proteins such as BRD2, BRD3 and BRD4 are epigenetic “readers” that interact with acetylated lysine residues in histones to regulate transcription. Principally based on the JQ1 compound^26^, a number of different BET PROTACs have been developed, including the VHL-recruiting ARV-771^27^ and MZ1^28^ tools, and the CRBN-recruiting dBET6^29^ compound.

As expected, ProtacID conducted in 293 cells with all three BET PROTACs identified BRD2, BRD3 and BRD4, along with a number of additional chromatin-associated proximity interactors that were also identified in a standard BioID analysis (selected high confidence hits presented in **Fig. 5**; all ProtacID data in **Supplementary Tables 9, 10** and see^30^). ProtacID conducted with the two VHL-recruiting BET PROTACs in two different cell lines (293 and 697) displayed 100% overlap within this conserved interactor set (**Supplementary Table 11**), and consistent with the presence of the same warhead, the CRBN-recruiting PROTAC dBET6 generated an analogous interactome in 293 cells (**Fig 5**).

**Figure 5.**
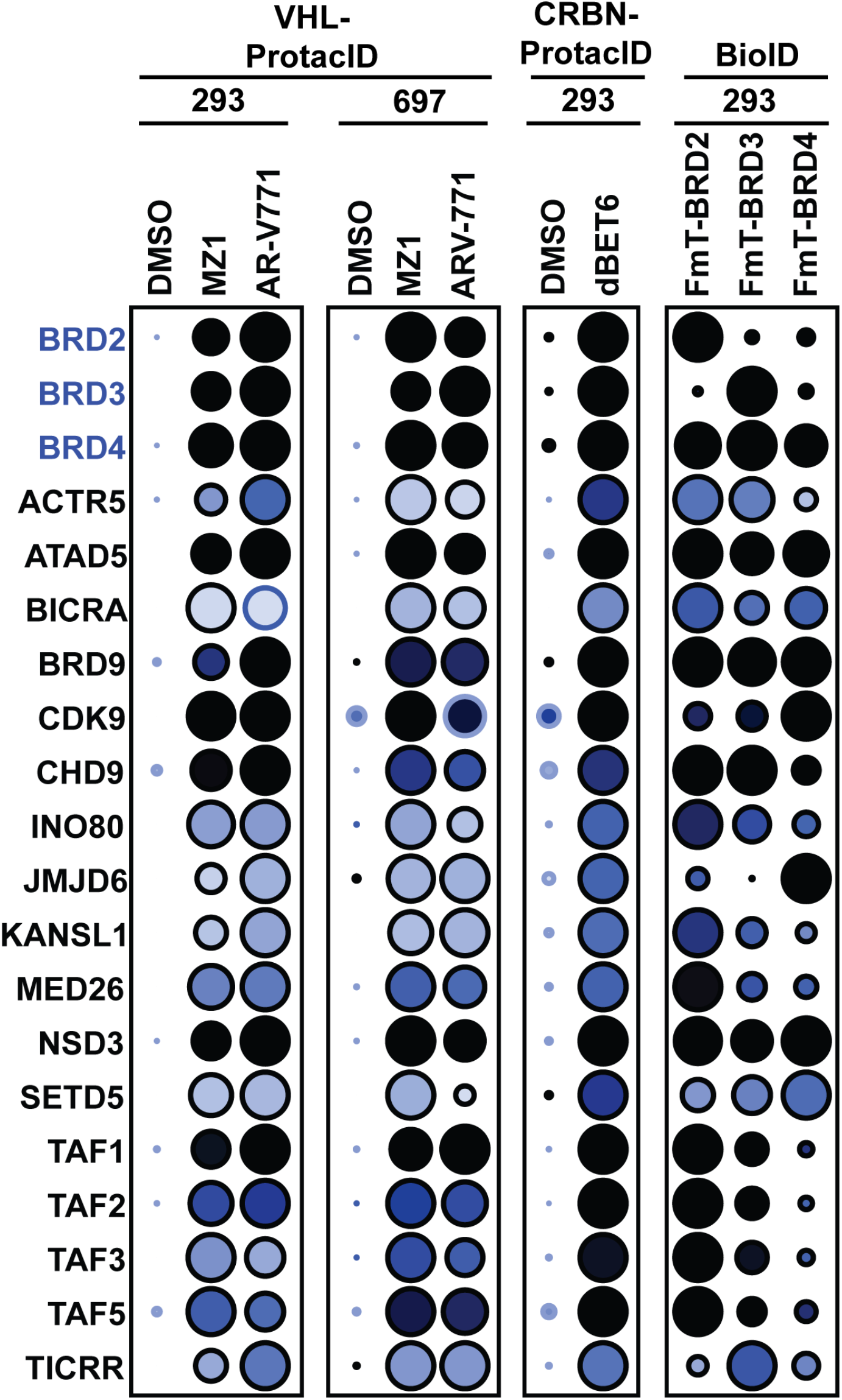
ProtacID conducted with three BET PROTACs. Two VHL-recruiting BET PROTACs (MZ1 and ARV-771), and one CRBN-recruiting BET PROTAC (dBET6) were subjected to ProtacID, as indicated. For comparison, standard BioID was also conducted on FmT-BRD2, FmT-BRD3 and FmT-BRD4 in 293 cells. *Blue* font indicates PROTAC neo-substrates.

### ProtacID characterization of EED-targeting PROTACs

The EED (embryonic ectoderm development) protein is a component of polycomb repressive complex 2 (PRC2), responsible for epigenetic silencing via methylation of histone H3 K27^31^. The PRC2 complex comprises the core subunits EED, EZH2 and SUZ12, along with additional variable accessory factors^32^. Two VHL-directed EED PROTACs, UNC7700^33^ and PROTAC2^34^, effect the degradation of EED, EZH2 and SUZ12 in the patient-derived non-Hodgkin lymphoma cell line Karpas-422. As expected, ProtacIDs conducted with EED PROTACs identified EED, EZH2 and SUZ12 as high confidence PROTAC proximity partners in Karpas-422 cells (**Fig 6a, Supplementary Table 11**). Consistent with previous AP-MS^35^ and BioID analyses^36^, the PRC2-associated AEBP2, MTF2 and PHF19 proteins were also identified in both ProtacID anlayses (**Fig 6a**).

**Figure 6.**
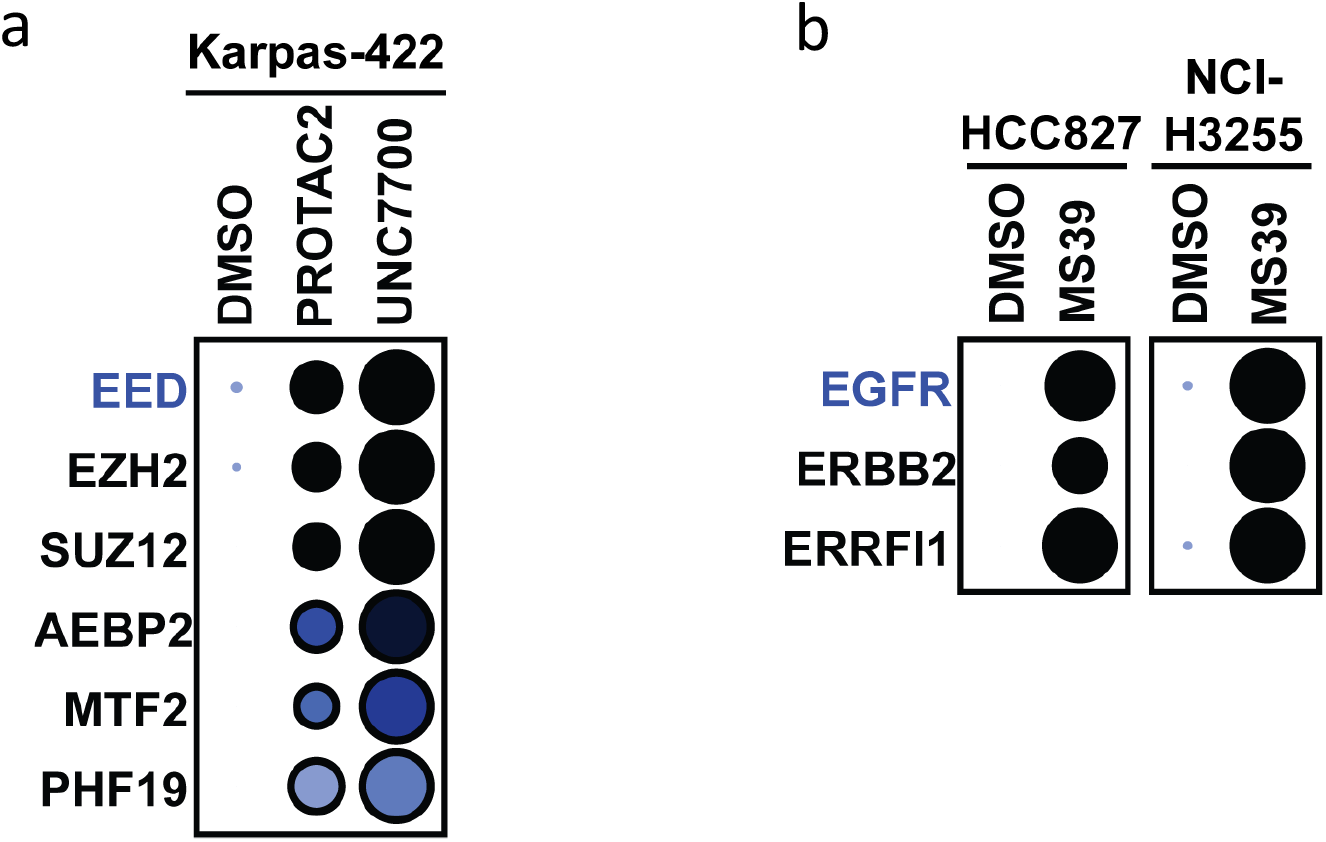
ProtacID conducted on (**a**) two EED-directed PROTACs in the Karpas-422 cell line, and (**b**) an EGFR PROTAC in the HCC827 and NCI-H3255 cell lines, as indicated. *Blue* font indicates PROTAC neosubstrates.

### ProtacID characterization of an EGFR PROTAC

Finally, it was important to determine whether the ProtacID approach can be applied to PROTACs that target proteins localized to other (non-chromatin) intracellular locations. The VHL-recruiting epidermal growth factor receptor (EGFR) PROTAC MS39^37^ (employing the tyrosine kinase inhibitor gefitinib^38^ as the warhead) was thus subjected to ProtacID. This PROTAC can target EGFR mutant proteins that drive non-small cell lung cancer and other diseases^39^. As expected, MS39 ProtacID identified the EGFR proteins expressed in the HCC827 (EGFR^Exdel19^) and NCI-H3225 (EGFR^L858R^) cell lines, as well as two previously reported membrane-localized EGFR interacting partners^40, 41^ (**Fig 6b, Supplementary Table 12**).

Taken together, our data demonstrate that ProtacID is a versatile tool that can be used to characterize VHL or CRBN-recruiting PROTACs directed against protein targets in different intracellular locations, and across different human cell types.

## Discussion

The development of TPD compounds has emerged as an important strategy to target proteins previously considered to be “undruggable” by small molecule inhibitors^1^. PROTACs thus hold great promise for the treatment of currently intractable diseases. However, as more PROTAC (and molecular glue)-based compounds progress to clinical use, it will become increasingly important to more fully characterize their interactomes *in vivo*.

The specificity of warhead compounds is ordinarily first characterized *in vitro* using recombinant target proteins or protein domains, and demonstrating that the compound of interest does not interact with (or displays significantly lower affinity for) related recombinant proteins in the same protein family. If the compound is used to create a PROTAC, the resulting tool is often then subjected to a standard global proteome analysis, which identifies proteins that change in abundance in response to the compound. However, this approach alone does not provide direct evidence for target engagement *in vivo*, nor can it identify non-productive interactions. Here we describe a simple strategy (**Fig 4e**) that can be used as a complementary approach to address this important need.

PROTAC development can also be slow and challenging. Even once highly specific warheads have been identified and optimized, combining them with standard linkers and E3 recruiter moieties does not always result in efficient target protein degradation (much less result in a compound with good “drug-like” properties). More rapid and straightforward methods for *in vivo* compound testing - such as ProtacID - could help to streamline this process, identifying compounds with promising, or unsatisfactory, specificities at earlier stages of development.

Future ProtacID development will focus on developing tools for the analysis of PROTACs and molecular glue compounds that recruit other ubiquitin E3 ligases, such as BIRC2, DCAF16, MDM2, XIAP and FBXO22^1, 2, 42-44^. And finally, while the quality of a ProtacID analysis will be dependent on the properties (*e*.*g*. solubility, cell permeability and specificity) of the associated PROTAC, we also report here that ProtacID can be used to generate highly accurate proximity interactome data similar to standard BioID approaches, identifying native, endogenous protein complexes without the use of antibodies, epitope tags or large biotin ligases fused to the protein of interest. This approach could be used to *e*.*g*. characterize how endogenous protein complexes are “re-wired” by specific coding mutations, or by other drugs or compounds, without any modification to the structure or expression level of the protein of interest.

## Methods

### Cloning

VHL and ΔVHL cDNAs were amplified by PCR from the VHL-pGex2TK plasmid (Addgene 20790). The nuclear localization sequence (NLS) PKKKRKVEDPKKKKKV was added to all VHL/ΔVHL constructs. To generate an inducible 293 Flp-In cell line, NLS-VHL/ΔVHL sequences were cloned into the pcDNA5 FRT/TO vector containing N-term and C-term 3xFLAG-miniTurbo using in-fusion HD EcoDry Mix (Takaro 639689). For the C-term miniTurbo variants, HindIII and NotI restriction sites were used. NsiI and EcoRV restriction sites were used to make N-term miniTurbo constructs. FLAG-miniTurbo-ΔVHL was additionally cloned into a modified pSTV6^45^ lentiviral vector using Xho1 and EcoR1 restriction sites. CRBN^midi^ (Addgene 215330) was cloned into pcDNA5 FRT/TO as above to generate the CRBN^midi^-miniTurbo-FLAG construct. A 12 amino acid linker (GGGGGSGGGGGS) was added between mini-turbo and CRBN^midi^ by PCR.

### Reagents

ACBI2 and VZ185 were obtained through the Boehringer Ingelheim opnMe program. UNC7700 was a kind gift from the L James laboratory (University of North Carolina, Chapel Hill, NC, USA). ACBI1 (catalog no. HY-128359, lot 66200), AU-15330 (HY-145388, lot 145137), AU-24118 (HY-163410, lot 535238), dBET6 (HY-112588, lot 113183), ARV-771 (HY-100972, lot 380611) and PROTAC 2, EED degrader (HY-130615, lot 113670) were purchased from MedChemExpress. MZ1 (catalog no. 21622, batch 0664999-3) and MLN4924 (15217, batch 0712600-6) were purchased from Cayman Chemical Company. MS39 (7397, batch 1A/293284) was purchased from Tocris Bioscience. All PROTACs were solubilized in DMSO and used as indicated.

### Cell culture

293 Flp-In T-REx cells were cultured in DMEM (Wisent) supplemented with 10% FBS (Wisent). 697 cells were cultured in RPMI medium (Wisent) supplemented with 10% heat-inactivated FBS (Gibco). The HAP1 parental cell line was a kind gift from the A Schimmer lab (Princess Margaret Cancer Centre). HAP1 KO lines (*SMARCA2KO* and *ARID1AKO*) were kind gifts from the S Kubicek lab (CeMM, The Research Center for Molecular Medicine, Vienna, Austria). HAP1 cells were cultured in IMDM (Iscove’s Modified Dulbecco Medium) with 10% FBS. The Karpas-422 cell line was a kind gift from the R Kridel lab (Princess Margaret Cancer Centre), and cultured in RPMI medium supplemented with 10% FBS. HCC827 and NCI-H3255 cell lines were kind gifts from the M Tsao lab (Princess Margaret Cancer Centre) and cultured in RPMI medium supplemented with 10% FBS. All media was supplemented with 100 U/ml penicillin and 100 ug/ml streptomycin (Wisent). Cells were cultured at 37°C in a humidified chamber with 5% CO_2_, following standard protocols, and tested for mycoplasma twice/year (MycoAlert Mycoplasma Detection kit, Lonza).

### Lentivirus production

One day before transfection, 0.5 million 293T cells were seeded in a 6-well plate. The next day, cells were transfected with FLAG-miniTurbo-ΔVHL pSTV6^45^ plasmid, along with the lentiviral transfer plasmid and the packaging plasmids pMDG VSVG and PAX2 (X-tremeGENE Millipore Sigma 6366244001) according to manufacturer’s instructions. Viral supernatants were collected 48h after transfection, combined and filtered through a 0.45 um Millex-HA syringe filter (Millipore Sigma SLHA033SS). Filtered supernatants were added to target cells in the presence of 8 ug/ml polybrene (Sigma-Aldrich TR-1003-G). After one day of incubation, viral supernatants were removed and replaced with normal media. Puromycin (2ug/ml) was added to the cells 48h post-infection to initiate selection for stably transduced cells.

### Generation of stable lines

293 Flp-In T-REx cells were co-transfected with pOG44 (Flp-recombinase expression vector) and pcDNA5-FRT/TO plasmid containing the coding sequences for FLAG-miniTurbo (FmT) fused in-frame with VHL, ΔVHL or CRBN^midi^. Transfections were performed using Lipofectamine 2000 (Invitrogen) according to manufacturer’s instructions. Following transfection, cells were selected with 200 ug/ml hygromycin. All other cell lines were transduced with the lentiviral vector encoding FLAG-miniTurbo-ΔVHL. 50-60% confluent cells in 6-well plates were infected with 350 µl of unconcentrated virus and polybrene to a final concentration of 8 µg/ml. Following transduction, cells were selected with 2 ug/ml puromycin.

### ProtacID

Cell lines stably expressing FmT-tagged VHL, DVHL or CRBN^midi^ proteins were divided into two treatment conditions, each comprising 5 × 15 cm^2^ plates at ∼80% confluency. Expression of the fusion protein was induced by treating the cells overnight with 2 ug/ml doxycycline. The following day, the cells were treated with 0.5 uM MLN4924 (Cayman Chemical 15217). After one hour, cells were treated with either PROTAC (based on DC_50_ values; ACBI1 0.2uM, ACBI2 0.1uM, AU-15330 0.4uM, VZ185 0.1uM, AU-24118 1uM, MZ1 0.5uM, ARV-771 0.2uM, dBET6 1uM, PROTAC2 1uM, UNC7700 1uM, MS39 1uM) or DMSO for three hours, followed by the addition of biotin to a final concentration of 50 uM for 60 min. At endpoint, culture medium was removed, cells were washed with chilled PBS and harvested by scraping. Cells were pelleted by centrifugation at 300×g for 5 min. The pellet was washed once with 10 ml PBS and centrifuged again at 300×g for at 5 min. PBS was removed, the cell pellet was flash-frozen in liquid nitrogen and stored at -80°C.

### BioID

BioID was conducted as previously described^46^. In brief, the cell pellet was resuspended in 10 ml lysis buffer (50 mM Tris-HCl pH 7.5, 150 mM NaCl, 1 mM EDTA, 1 mM EGTA, 1% Triton X-100, 0.1% SDS, 0.5% sodium deoxycholate, 1:500 protease inhibitor cocktail (Sigma-Aldrich), 1:1,000 benzonase nuclease (Novagen 71205-M) and incubated on an end-over-end rotator at 4°C for 1 h. The lysate was briefly sonicated to disrupt any visible aggregates, then centrifuged at 45,000×g for 30 min at 4°C. Supernatant was transferred to a fresh 15 ml conical tube. 30ul of packed, pre-equilibrated streptavidin-sepharose beads (Cytiva 17-5113-01) were added, and the mixture incubated for 3h at 4°C with end-over-end rotation. Beads were pelleted by centrifugation at 376×g for 2 min and transferred with 1 ml of lysis buffer to a fresh Eppendorf tube.

An aliquot (∼10%) of the beads was saved for immunoblotting. The remainder of the beads was washed four times with 50 mM ammonium bicarbonate (ammbic, pH 8.3) and transferred to a fresh centrifuge tube for two additional washes with 1 ml of ammbic. Tryptic digestion was performed by incubating the beads with 1ug MS-grade TPCK trypsin (Promega V5280) in 200ul 50 mM ammbic (pH 8.3) overnight at 37°C. The following morning, an additional 0.5 ug trypsin was added, and the beads incubated for another 2h at 37°C. Beads were then pelleted by centrifugation at 2,000×g for 2 min, and the supernatant transferred to a fresh Eppendorf tube. Beads were washed twice with 200 ul 50 mM ammbic, and the washes pooled with the eluate. The sample was lyophilized and resuspended in buffer A (0.1% formic acid). One-fifth of each sample was analyzed per MS run.

### Mass Spectrometry (**ProtacID**)

High performance liquid chromatography (HPLC) was conducted using a 2 cm pre-column (Acclaim PepMap 50 mm x 100 um inner diameter) and 50 cm analytical column (Acclaim PepMap, 500 mm x 75 um diameter; C18; 2 um; 100 Å, Thermo Fisher Scientific, Waltham, MA) running a 120 min reversed-phase buffer gradient at 225 nl/min on a Proxeon EASY-nLC 1000 pump in-line with a Thermo Q-Exactive HF quadrupole-Orbitrap mass spectrometer. A parent ion scan was performed using a resolving power of 60,000, then up to the twenty most intense peaks were selected for MS/MS (minimum ion count of 1,000 for activation) using higher energy collision induced dissociation (HCD) fragmentation. Dynamic exclusion was activated such that MS/MS of the same *m*/*z* (within a range of 10 ppm; exclusion list size = 500) detected twice within 5 sec were excluded from analysis for 15 sec.

For protein identification, Thermo .RAW files were converted to the .mzXML format using Proteowizard^47^, then searched using X!Tandem^48^ and COMET^49^ against Human RefSeq Version 45 database (containing 36,113 entries). Data were analyzed using the trans-proteomic pipeline (TPP)^50^ via the ProHits software suite (v3.3)^51^. Search parameters specified a parent ion mass tolerance of 10 ppm, and an MS/MS fragment ion tolerance of 0.4 Da, with up to 2 missed cleavages allowed for trypsin. Variable modifications of +16@M and W, +32@M and W, +42@N-terminus, and +1@N and Q were allowed. Proteins identified with an iProphet cut-off of 0.9 (corresponding to ≤1% FDR) and at least two unique peptides were analyzed with SAINT Express v.3.6.13^52^. Twenty control runs (from cells stably expressing the FmT epitope tag only) were collapsed to the four highest spectral counts for each prey and compared to two biological replicates (each with two technical replicates). High confidence interactors were defined as those with a Bayesian false discovery rate (BFDR) ≤0.01. High confidence PROTAC-mediated interactions were defined as those with >5 average peptide counts/MS run and >2-fold (log_2_) spectral counts in the presence *vs* absence of PROTAC.

### Global Proteome Analysis

Cell pellets were resuspended in 100ul lysis buffer (5% SDS, 50 mM triethylammonium bicarbonate pH 8.5) and sonicated. Following centrifugation, cleared lysates were reduced (5 mM DTT 60°C 30min), alkylated (20 mM iodoacetamide RT 15min), then acidified to a final concentration of 1.2% H_3_PO_4_. Lysates were diluted (1/7) with 100 mM TEAB/90% methanol, and 192.5 ul were loaded onto S-trap™ micro columns (ProtiFi, Fairport, NY) followed by washing according to manufacturer’s instructions. On-column digest was conducted with 2ug MS-grade TPCK trypsin (Promega V5280) overnight at 37°C. Peptides were eluted from the S-Trap (50 mM TEAB, 0.2% HCOOH and CH_3_CN, successively) and desalted, and 500ng of sample was loaded onto an Evotip Pure™ column (Evosep EV2011). Liquid chromatography was performed with the Evosep One (Evosep) pump with an SPD30 method using a PepSep™ C18 HPLC column (15 cm x 150 µm ID, 1.5 µm; Bruker, Bremen). The TIMS-TOF HT (Bruker, Bremen) mass spectrometer was operated in PASEF-DIA^20^ positive ion mode (MS scan range 100-1700 *m/z*). Ion mobility range was 1/K0 = 1.6 to 0.6 Vs cm-2 using equal ion accumulation and ramp time in the dual TIMS analyzer of 100 ms each. DIA (scan range 400-1201 *m/z*) mass window width was 26 Da with 1 Da overlap and 32 steps per cycle. Collision energy was lowered stepwise as a function of increasing ion mobility, starting from 20 eV for 1/K0 = 0.6 Vs cm-2 and 59 eV for 1/K0 = 1.6 Vs cm-2. The ion mobility dimension was calibrated linearly using three ions from the Agilent ESI LC/MS tuning mix (*m/z*, 1/K0: 622.0289, 0.9848 Vs cm-2; 922.0097, 1.1895 Vs cm-2; and 1221.9906, 1.3820 Vs cm-2). Data files were analyzed on the Fragpipe platform^21^ (v20.0) using DIA-NN^22^ (v1.8.2_beta_8) with a Bruker spectral library generated from K562 and Molt4 trypsin-digested cell lysates fractionated using high pH reversed-phase chromatography and analyzed on a TIMS-TOF HT.

### Global Proteome Data Analysis

Protein groups were loaded into R (4.3.1, R Core Team (2020), https://www.R-project.org/). Proteins labeled as known contaminants in the spectral library and proteins not quantified in all replicates of at least one condition were removed from analysis. Imputation of missing data (random draws from a manually defined left-shifted Gaussian distribution (shift = 1.8, scale = 0.3)) and testing for protein differential abundance was done using the R package differential enrichment analysis of proteomics data (DEP) (1.24.0) as described in the package documentation.

### Western blotting

ProtacID blots. Aliquots (∼10%) of protein-bound beads from ProtacID samples were washed 4x with cell lysis buffer, heated in 50ul 4×NuPAGE buffer at 95°C for 5 min, then centrifuged in a tabletop centrifuge at max speed for 10 min. The supernatant was transferred to a fresh Eppendorf tube for immunoblotting analysis.

For blotting of whole cell lysates, cells were collected as above, and pellets lysed for 3 min RT in 20 mM Tris-HCl (pH 8), 150 mM NaCl, 10 mM MgCl_2_, 1 mM EDTA, 0.5% Triton X-100, protease inhibitor (Roche 11873580001) and benzonase (Millipore Sigma E1014). SDS was added to a final concentration of 1%. Lysates were then spun in a tabletop centrifuge at max speed for 10 min, and supernatants transferred to fresh tubes. Protein concentrations were measured (Thermo Scientific Pierce™ BCA Protein Assay Kit 23225) according to manufacturer protocol. 10-30 µg total protein was loaded into Invitrogen NuPAGE Bis-Tris Gels and run in 1X NuPAGE MOPS buffer (Invitrogen NP0001) for 90 min at 120V. Proteins were transferred to 0.2 um polyvinylidene difluoride (PVDF, Cytiva) membrane using a Bio-Rad Mini Trans-Blot Cell at 70 V/500 mA for 1.5 hr in Tris-Glycine transfer buffer (3 g/L Tris, 14.4 g/L glycine, and 10% methanol) on ice. Membranes were blocked in 5% BSA in PBS-T for 1 hr at room temperature, followed by overnight incubation with primary antibodies: FLAG (1:5000 Sigma-Aldrich, F1804, Mouse), SMARCA4 (BRG1 1:1000; Cell Signaling 49360T, Rabbit), and MPHOSPH1 (KIF20B 1:1000; Invitrogen PA5-120351, Rabbit) diluted in 5% BSA. After primary antibody incubation, membranes were washed 3 × 10 min with PBS-T, then incubated for 1h at RT with LI-COR secondary antibodies: IRDye 800CW Goat anti-Rabbit (LI-COR 926-32211), or 680RD Goat anti-Mouse (926-68070), all diluted 1:5000 in 5% BSA. Membranes were washed 3 × 10 min with PBS-T then imaged using a LI-COR Odyssey CLx scanner.

## Supplementary Figure Legends

**Supplementary Figure 1.**
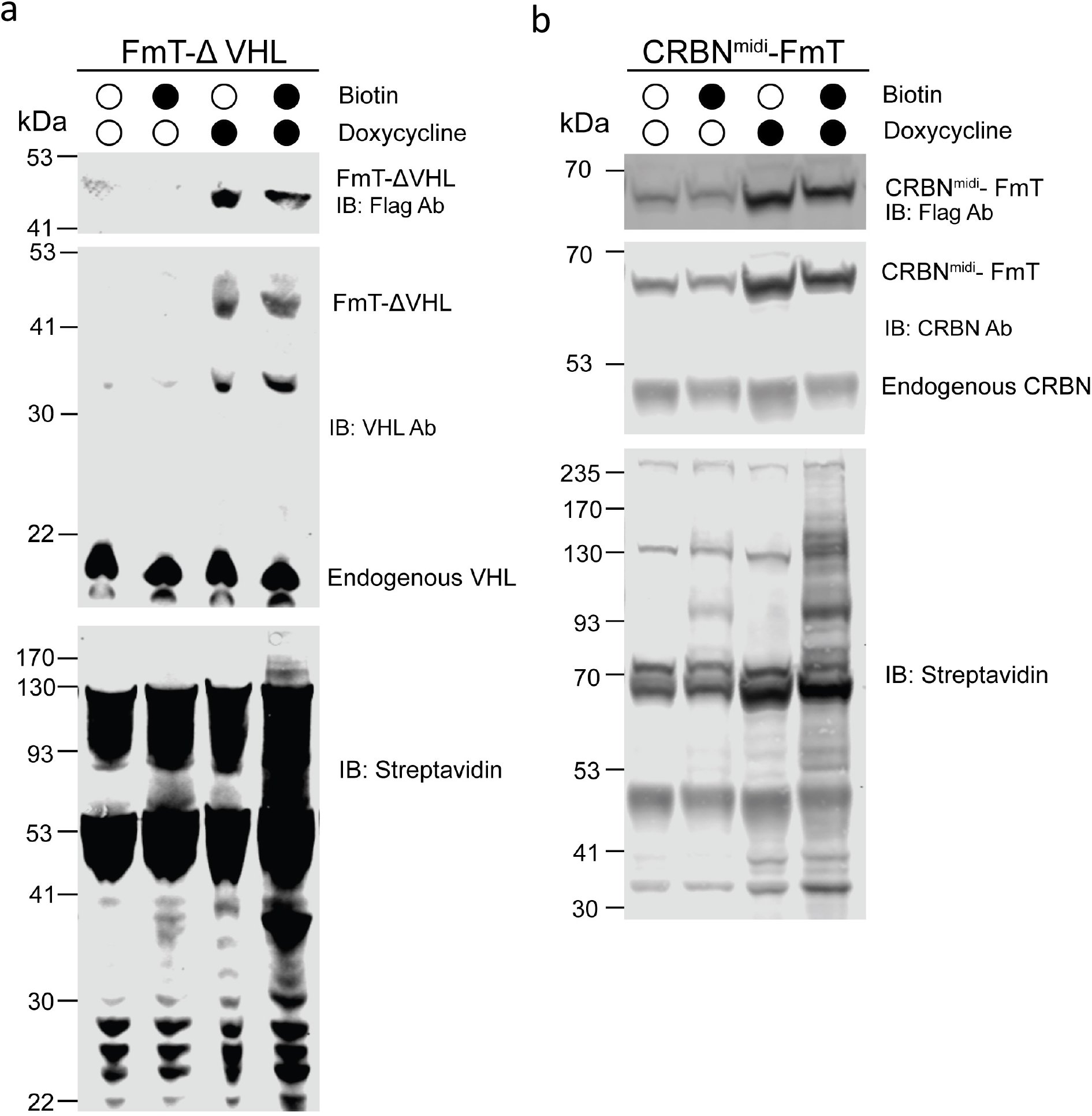
293 Flp-In cells expressing (**a**) FmT-ΔVHL or (**b**) CRBN^midi^-mTF were treated as indicated, and whole cell lysates subjected to western blotting. Doxycycline - 2ug/ml added to culture media for 24 hrs. Biotin - 50uM biotin for 1hr.

**Supplementary Figure 2.**
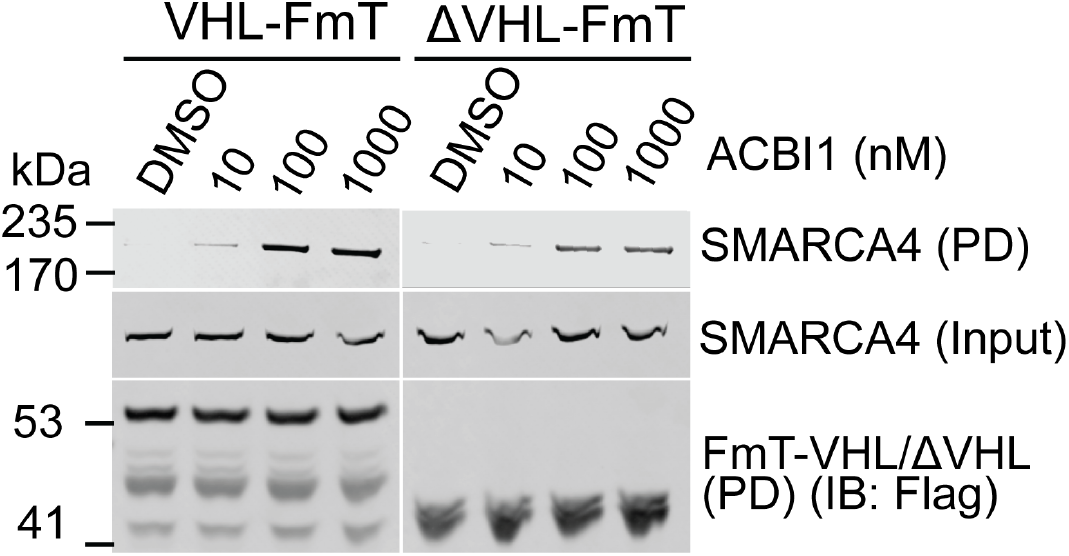
Increasing doses of ACBI1 were applied to 293 Flp-In cells expressing the indicated C-terminally FmT-tagged VHL/ΔVHL proteins. After one hour of biotin treatment, cells were lysed and biotinylated proteins isolated using streptavidin-sepharose (*PD*, pulldown). Isolated proteins (or whole cell lysates, *input*) were subjected to Western blotting with the indicated antibodies.

## Supplementary Tables

1. ACBI1 ProtacID conducted in 293 Flp-In cells with four different VHL protein variants (full-length or ΔVHL, tagged at the N- or C-terminus with FLAG-miniTurbo).
2. Standard BioID conducted on FmT-SMARCA2 and FmT-SMARCA4 in 293 Flp-In cells.
3. FmT-ΔVHL ProtacID conducted in 293 Flp-In cells treated with four different BAF PROTACs: ACBI1, ACBI2, AU-15330 and VZ185.
4. FmT-ΔVHL ProtacID conducted in 697 cells with the BAF PROTACs ACBI1 and ACBI2.
5. FmT-ΔVHL ProtacID conducted in WT, *ARID2* KO and *SMARCA2* KO HAP1 cells treated with ACBI1.
6. CRBN^midi^-mTF ProtacID conducted in 293 Flp-In cells treated with the BAF PROTAC AU-24118.
7. Global Proteome analysis of 293 Flp-In cells treated with ACBI1.
8. FmT-ΔVHL ProtacID conducted in 293 Flp-In cells treated with ACBI1, in the absence of MLN4924.
9. FmT-ΔVHL ProtacID conducted in 293 Flp-In cells treated with the BET PROTACs ARV-771 or MZ1.
10. CRBN^midi^-mTF ProtacID conducted in 293 Flp-In cells treated with the BET PROTAC dBET6.
11. FmT-ΔVHL ProtacID conducted in 697 cells treated with the BET PROTACs ARV-771 or MZ1.
12. Standard BioID conducted on FmT-BRD2, FmT-BRD3 and FmT-BRD4 in 293 Flp-In cells.
13. FmT-ΔVHL ProtacID conducted in Karpas-422 cells treated with the EED PROTACs UNC7700 and AZ EED PROTAC2.
14. FmT-ΔVHL ProtacID conducted in HCC827 cells treated with the EGFR PROTAC MS39.
15. FmT-ΔVHL ProtacID conducted in NCI-H3255 cells treated with EGFR PROTAC MS39.

## Data availability

All .raw mass spectrometry files are publicly available at the MassIVE data repository (massive.ucsd.edu), ID MSV000095096. Reviewer password: ProtacID_2024.

## Biological Material availability

Plasmids will be made available at Addgene upon manuscript acceptance.

### Acknowledgements

We thank the Gingras lab for the pDEST-pcDNA5 vector.

SS was supported by a Princess Margaret Cancer Centre post-doctoral fellowship. MERM is supported by a Canadian Institutes of Health Research (CIHR) post-doctoral fellowship.

LJ is supported by a Mitacs Accelerate Award.

DYN was supported by a Canada Graduate Scholarship – Doctoral Research Award from CIHR (494204) and a Doctoral Training Scholarship from Fonds de recherche du Québec - Santé (320128).

JS-G is supported by generous funding from The Princess Margaret Cancer Centre Foundation. CHA was supported by funding from CIHR grants FDN154328 & OGB190363. The Structural Genomics Consortium is a registered charity (no: 1097737) that receives funds from Bayer AG, Boehringer Ingelheim, Bristol Myers Squibb, Genentech, Genome Canada through The Ontario Genomics Institute [OGI-196], EU/EFPIA/OICR/McGill/KTH/Diamond Innovative Medicines Initiative 2 Joint Undertaking [EUbOPEN grant 875510], Janssen, Merck KGaA (aka EMD in Canada and US), Pfizer and Takeda.

BR holds the Philip S. Orsino Chair in Leukemia Research, administered through the University of Toronto and The Princess Margaret Cancer Centre. Work in the BR lab was supported by the CIHR project grant program, and additional generous funding from The Princess Margaret Cancer Centre Foundation.

## Author contributions

Conceptualization: SS, MERM, DYN, DB-L, CHA, BR. Investigation: SS performed all VHL ProtacID experiments. MERM performed CRBNmidi ProtacID experiments and data analysis for global proteomics. LJ performed all standard BioID experiments. SD conducted cloning and cell culture work. DYN generated cell lines, conducted western blotting. JS-G performed all mass spectrometry and MS data analysis. SS, CHA and BR wrote the paper. All authors contributed to the manuscript.

